# Association of *CXCR6* with COVID-19 severity: Delineating the host genetic factors in transcriptomic regulation

**DOI:** 10.1101/2021.02.17.431554

**Authors:** Yulin Dai, Junke Wang, Hyun-Hwan Jeong, Wenhao Chen, Peilin Jia, Zhongming Zhao

## Abstract

**Background:** The coronavirus disease 2019 (COVID-19) is an infectious disease that mainly affects the host respiratory system with ∼80% asymptomatic or mild cases and ∼5% severe cases. Recent genome-wide association studies (GWAS) have identified several genetic loci associated with the severe COVID-19 symptoms. Delineating the genetic variants and genes is important for better understanding its biological mechanisms.

**Methods:** We implemented integrative approaches, including transcriptome-wide association studies (TWAS), colocalization analysis and functional element prediction analysis, to interpret the genetic risks using two independent GWAS datasets in lung and immune cells. To understand the context-specific molecular alteration, we further performed deep learning-based single cell transcriptomic analyses on a bronchoalveolar lavage fluid (BALF) dataset from moderate and severe COVID-19 patients.

**Results:** We discovered and replicated the genetically regulated expression of *CXCR6* and *CCR9* genes. These two genes have a protective effect on the lung and a risk effect on whole blood, respectively. The colocalization analysis of GWAS and *cis*-expression quantitative trait loci highlighted the regulatory effect on *CXCR6* expression in lung and immune cells. In the lung resident memory CD8^+^ T (T_RM_) cells, we found a 3.32-fold decrease of cell proportion and lower expression of *CXCR6* in the severe than moderate patients using the BALF transcriptomic dataset. Pro-inflammatory transcriptional programs were highlighted in T_RM_ cells trajectory from moderate to severe patients.

**Conclusions:** *CXCR6* from the *3p21*.*31* locus is associated with severe COVID-19. *CXCR6* tends to have a lower expression in lung T_RM_ cells of severe patients, which aligns with the protective effect of *CXCR6* from TWAS analysis. We illustrate one potential mechanism of host genetic factor impacting the severity of COVID-19 through regulating the expression of *CXCR6* and T_RM_ cell proportion and stability. Our results shed light on potential therapeutic targets for severe COVID-19.

## Background

The coronavirus disease 2019 (COVID-19) pandemic has already infected over 100 million people and caused numerous morbidities and over 2 million death worldwide as of January 2021. The virus is evolving fast with new variants being emerged in the world [1, 2]. A huge disparity in the severity of symptoms in different patients has been observed. In some of the patients, only mild symptoms or even no symptoms are shown and little treatment or interventions are required while a subset of patients experience rapid disease progression to respiratory failure and need urgent and intensive care [3]. Although age and sex are major risk factors of COVID-19 disease severity [4], it remains largely unclear about the factors leading to the variability on COVID-19 severity and which group of individuals confer intrinsic susceptibility to COVID-19.

Several genome-wide association studies (GWAS) have been carried out and one genomic risk locus, *3p21*.*31*, has been replicated to be associated with the critical illness. One recent study by the Severe COVID-19 GWAS Group identified *3p21*.*31* risk locus for the susceptibility to severe COVID-19 with respiratory failure [5]. This GWAS signal was then replicated in a separate meta-analysis comprising in total 2,972 cases from 9 cohorts by COVID-19 Host Genetics Initiative (HGI) round 4 alpha. However, there is a cluster of 6 genes (*SLC6A20, LZTFL1, CCR9, FYCO1, CXCR6*, and *XCR1*) nearby the lead SNP rs35081325 within a complex linkage disequilibrium (LD) structure, which makes the “causal” gene and functional implication of this locus remain elusive [5, 6].

The majority of GWAS variants are located in non-coding loci, many of which are in the enhancer or promoter regions, playing roles as *cis*- or *trans*-regulatory elements to alter gene expression [7]. Although the function of non-coding variants could not be directly interrupted by their locations, their mediation effect on gene expression could be inferred by the expression quantitative trait loci (eQTL) analysis. In recent years, large consortia like GTEx (Genotype-Tissue Expression), eQTLGen Consortium, and DICE (database of immune cell expression) have generated rich eQTLs resources in diverse tissues and immune-related cell types [7-9]. A variety of statistical approaches such as transcriptome-wide association study (TWAS) analysis and colocalization analysis have successfully interpreted the target genes of non-coding variants by integrating the context-specific eQTLs [10-13].

Recent advances in single cell transcriptome sequencing provide unprecedented opportunities to understand the biological mechanism underlying disease pathogenesis at the single cell and cell type levels [14-16]. The recent generation of single cell RNA-sequencing (scRNA-seq) data from the bronchoalveolar lavage fluid (BALF) of moderate and severe COVID-19 patients has revealed the landscape of the gene expression changes in major immune cells. However, the transcriptome alteration in specific subpopulations remains mostly unexplored [17].

In this study, we aimed to connect the genetic factors with the context-specific molecular phenotype in COVID-19 patients. As illustrated in **Fig. 1**, we designed a multi-level workflow to dissect the genetically regulated expression (GReX) that contributed to severe COVID-19. We performed TWAS and colocalization analyses with a broad collection of eQTL datasets at the tissue and cellular levels. We further integrated the BALF single cell transcriptome dataset to explore the cellular transcriptome alterations in severe and moderate COVID-19 patients. Lastly, we proposed a hypothetical mechanism, connecting our multi-layer evidence in host genetic factors, gene (*CXCR6*), and single cell transcriptome features with the severity of COVID-19.

**Fig 1.**
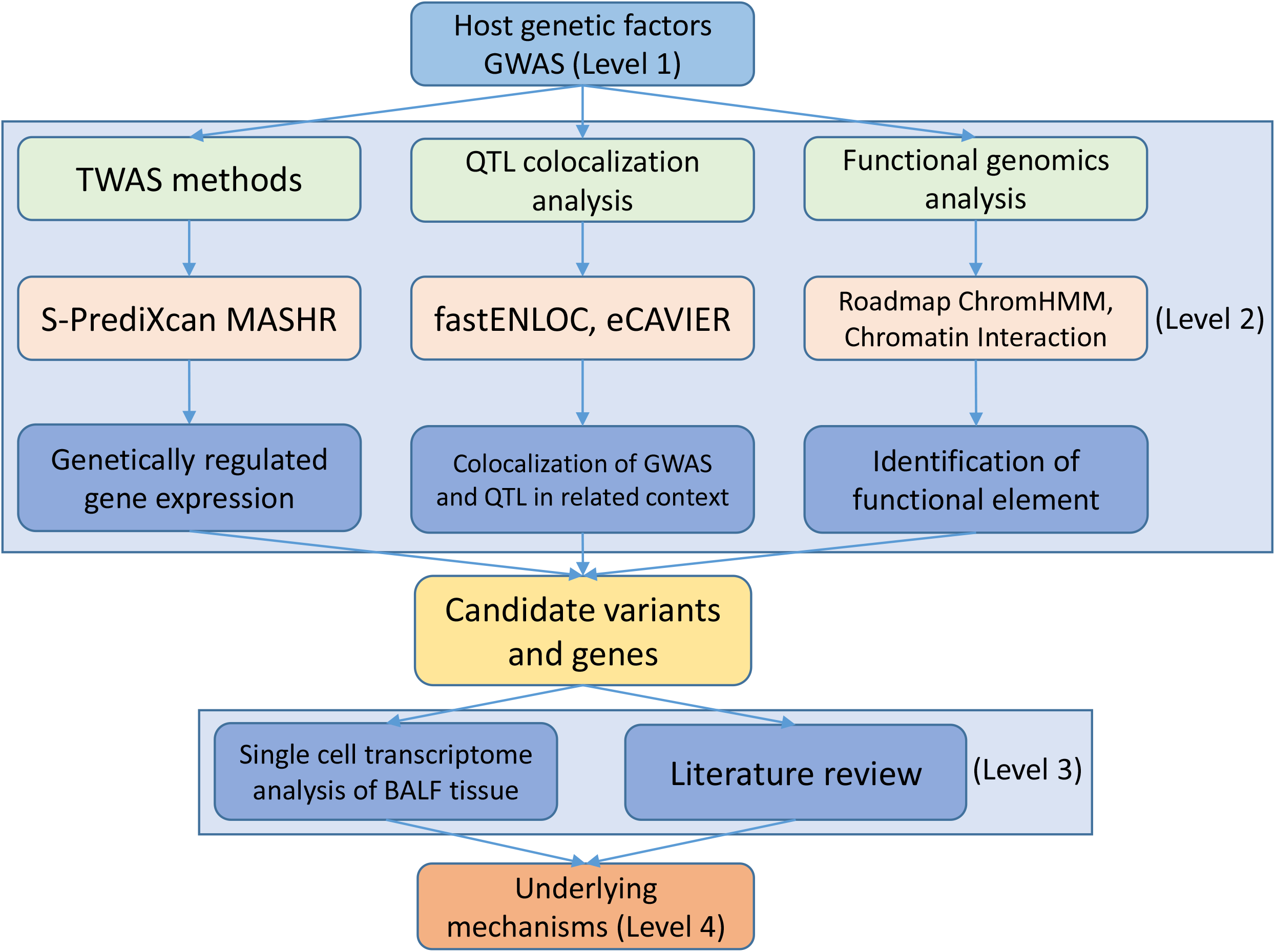
Workflow of a data-driven study: from genetic factor to molecular phenotype. The study has four major levels. Level 1: we collected the current largest COVID-19 genome-wide association study (GWAS) datasets and a non-duplicated replicate of the severe COVID-19 GWAS dataset. Level 2: we utilized the cutting-edge statistical approaches (transcriptome-wide association study and colocalization analysis) and public functional genomics annotations to dissect the genetic effects on gene expression (Methods). Then, we cross-validated our findings of these methods to ensure the robustness. Level 3: we adapted single cell RNA sequencing dataset from COVID-19 bronchoalveolar lavage fluid samples. We applied differentially expressed gene analysis and machine learning methods to characterize the molecular changes of candidate gene at single cell level from COVID-19 moderate and severe patients. We conducted extensive literature review to explain our observations. Level 4: we proposed a mechanism for explaining the “causal” association of genetic factors and the severity of COVID-19 patients.

## Methods

### GWAS dataset

We obtained GWAS summary statistics for the phenotype “severe COVID-19 patients vs population” (severe COVID-19) from two separate meta-analyses carried out by the COVID-19 Host Genetics Initiative (HGI, https://www.covid19hg.org/) and the Severe COVID-19 GWAS Group (SCGG) [5]. The GWAS_HGI_ A2 round 4 (alpha) cohort consists of 12,816,037 SNPs from the association study of 2,972 very severe respiratory confirmed COVID-19 cases and 284,472 controls with unknown SARS-CoV-2 infection status from nine independent studies in a majority of the European Ancestry population. The GWAS_SCGG_ dataset is from the first GWAS of severe COVID-19 [5], including 8,431,427 SNPs from the association study conducted from 1,980 COVID-19 confirmed patients with severe disease status and 2,205 control participants from two separate cohorts in Europe.

### Transcriptome-wide association analysis

We performed TWAS analyses of severe COVID-19 using S-PrediXcan [18] to prioritize GWAS findings and identify eQTL-linked genes. S-PrediXcan is a systematic approach that integrates GWAS summary statistics with publicly available eQTL data to translate the evidence of association with a phenotype from the SNP level to the gene level. Briefly, prediction models were built by a flexible and generic approach multivariate adaptive shrinkage in R package (MASHR) using variants with a high probability of being causal for QTL and tissue expression profiles from the GTEx version 8 [7, 19]. We chose three tissues that were relevant to SARS-CoV-2 infection, including lung, whole blood, and spleen. Then, we ran S-PrediXcan scripts (downloaded from https://github.com/hakyimlab/MetaXcan, accessed on 10/10/2020) with each of the three tissue-specific models in two severe COVID-19 GWAS datasets respectively. The threshold used in TWAS significance was adjusted by Bonferroni multiple test correction with the ∼10,000 genes. We defined the strict significance as p < 5 × 10^−6^ (|z| > 4.56) and suggestive significance as p < 5 × 10^−5^ (|z| > 4.06).

### Colocalization analysis

Colocalization was performed to validate significant TWAS associations using two recent and cutting-edge statistical analysis approaches: eCAVIAR [20] and fastENLOC [21], which aim to identify a single genetic variant that has shared causality between expression and GWAS trait. Both eCAVIAR and fastENLOC could assess the colocalization posterior probability (CLPP) for two traits at a locus, while eCAVIAR allows for multiple causal variants and fastENLOC features accountability for allelic heterogeneity in expression traits and high sensitivity of the methodology. We ran eCAVIAR between significant TWAS genes and GWAS trait with a maximum of five causal variants per locus and defined a locus as 50 SNPs up- and down-stream of the tested causal variant, following the recommendation in the original paper. The eCAVIAR was downloaded from https://github.com/fhormoz/caviar/ (accessed on 10/25/2020). The biallelic variants from the 1,000 Genomes Project phase III in European ancestry were used as an LD reference [22]. We defined CLPP > 0.5 as having strong colocalization evidence.

To run fastENLOC, we first prepared probabilistic eQTL annotations to generate the cis-eQTL’s posterior inclusion probability (PIP). Specifically, we applied the tissue-specific data from GTEx and T follicular cell-specific data from the DICE database [9] using the integrative genetic association analysis with the deterministic approximation of posteriors (DAP-G) package [23]. Then, GWAS summary statistics were split into approximately LD-independent regions defined by reference panel from European ancestry and z-scores were converted to PIP. We downloaded the fastENLOC from https://github.com/xqwen/fastenloc (accessed on 10/25/2020) and followed the guideline to yield regional colocalization probability (RCP) for each independent GWAS locus using each tissue-or cell type-specific eQTL annotation. We defined RCP > 0.5 as having strong colocalization evidence.

### Functional genomics annotations

To better understand the potential function of the variants identified by GWAS analyses and how they mediate the regulatory effect, we annotated significant SNPs using publicly available data. We obtained the tissue and cellular level eQTL data from the following resources: 1) the eQTLGen consortium [24] eQTLs generated from 30,912 whole blood samples; 2) Biobank-based Integrative Omics Studies (BIOS) eQTLs generated from 2,116 healthy adults [25]; 3) The GTEx v8 [7] eQTLs of the lung, whole blood, and spleen tissues; 4) DICE database [9] with cellular eQTLs of 9 available T cell subpopulations. To identify the genomic annotation of the significant SNPs, we downloaded the multivariate hidden Markov model (ChromHMM) [26] processed chromatin-state data of 17 lung and T cell lines from the Roadmap Epigenomics project [27]. To explore the potential chromatin looping of GWAS locus, we used publicly available chromatin interaction (Hi-C) data [28] at a resolution of 40Kb on IMR90, a normal lung fibroblast cell line. The Hi-C data has been used to identify specific baits and targets from distant chromatin regions that frequently interact with each other. Variants within the regulatory regions can be connected to the potential gene targets and thus mediate the gene expression. Statistical tests of bait-target pairs were conducted to define significant bait interaction regions and their targets. The eQTL associations and chromatin-state information and Hi-C interactions were processed and plotted using the R Bioconductor package gviz in R version 4.0.3 [29].

### Differentially expressed gene analysis in resident memory CD8^+^ T cells

We use the recently published scRNA-seq dataset of bronchoalveolar lavage fluids (BALF) samples from nine patients (three moderate and six severe) with COVID-19 [17, 30]. We adapted the original annotation [17] and followed their method to calculate the resident memory CD8^+^ T (T_RM_) cells signature score by using 31 markers (14 positive markers and 17 negative markers) for all annotated CD8^+^ T cells [31, 32]. We excluded cells with CD4^+^ expression and defined the top 50% scored cells as the T_RM_ cells. Lastly, we conducted a non-parametric Wilcoxon rank sum test by the function of “FindAllMarkers” from R package Seurat [33](version 3.1.5 in R version 3.5.2) to perform the differentially expressed genes (DEG) analysis between moderate and severe patients.

### Cell trajectory and transcriptional program analysis in T_RM_ cells

We used the R package Slingshot [34] to infer cell transition and pseudotime from the scRNA-seq data. Specifically, we first used the expression data to generate the minimum spanning tree of cells in a reduced-dimensionality space [t-Distributed Stochastic Neighbor Embedding (tSNE) project from top 30 principle components of top 3,000 variable genes] assuming there are two major clusters (moderate and severe T_RM_ cells). We then applied the principal curve algorithm [35] to infer an one-dimensional variable (pseudotime) representing the each cell’s trajectory along the transcriptional progression. We used our in-house machine learning tool, DrivAER (Driving transcriptional programs based on AutoEncoder derived relevance scores) [36], to identify potential transcriptional programs (e.g., gene sets of pathways or transcription factors (TF)s) that potentially regulate the inferred cell trajectory between the moderate and severe patients. To avoid the potential noise from the low expression genes, we excluded those genes expressed in < 10% cells. DrivAER took gene-expression and pseudotime inferred from previous cell trajectory results (Slingshot) and calculated each gene’s relevance score by performing cellular manifold by using Deep Count Autoencoder [37] and a random forest model with out-of-bag score calculation as the relevance score. The transcriptional program annotations were from the hallmark pathway gene sets from MSigDB [38] and transcription factor (TF) target gene sets from TRRUST [39]. To calculate the relevance score, we used the “calc_relevance” function with the following parameters: min_targets = 10, ae_type = “nb-conddisp”, epoch=100, early_stop=3, and hidden_size = “(8,2,8)”. The relevance score (R^2^ coefficient of determination) indicates the proportion of variance in the pseudotime explained by target genes of transcription factor or genes in the hallmark pathways.

### DNA motif recognition analysis of genome-wide significant SNPs

We used the function “variation-scan” of the online tool RSAT (http://rsat.sb-roscoff.fr/index.php, accessed on 01/15/2020) [40] to predict the binding effect of all the significant SNPs at the *3p21*.*31* locus. We defined the TF with Bonferroni corrected p < 0.05 as the significant TF. Later, we compared them with the TF with high relevance score from the DrivAER analysis above. The position weight matrices (PWMs) for all the TFs were downloaded from cis-BP Database (http://cisbp.ccbr.utoronto.ca/) version 2019-06_v2.00) [41] and sequence logos representing motif binding sites were generated using R package seqLogo version 1.54.3 in R version 3.5.2.

## Results

### TWAS analysis identified and replicated two chemokine receptor genes

We utilized the latest S-PrediXcan MASHR models trained with GTEx v8 data for TWAS analyses in lung and whole blood on two GWAS datasets of susceptibility to severe COVID-19 [19]. In the HGI cohort, we found that a decreased expression of *CXCR6*, which encodes C-X-C chemokine receptor type 6, in the lung was associated with an increased risk for the development of severe COVID-19 symptoms (p = 1.57 × 10^−17^, z = -8.53), and this result was then replicated in the SCGG cohort (p = 2.84 × 10^−5^, z = -4.19, suggestive significant) (**Fig. 2** and **Table 1**). Likewise, an increased expression of *CCR9*, which encodes C-C chemokine receptor type 9, in whole blood was associated with an increased risk for the development of severe COVID-19 complications in GWAS_HGI_ cohort (p = 7.90 × 10^−11^, z = 6.50) and this result was replicated in the other GWAS_SCGG_ cohort, (p = 3.78 × 10^−10^, z = 6.26) (**Fig. 2** and **Table 1**). Whole blood and lung transcriptome models also identified two additional significant TWAS genes that are specific to one of the two cohorts. Increased expression of *ABO* gene in the lung was associated with risk for the development of severe COVID-19 symptoms in GWAS_SCGG_ data set (p = 5.98 ×10^−7^, z = 4.99). Similarly, increased expression of *GAS7* gene (Growth Arrest-Specific 7) in whole blood was associated with an increased risk for development of COVID-19 symptom in the GWAS_HGI_ data set (p = 8.46 × 10^−7^, z = 4.92). Overall, these two chemokine receptor genes were found and replicated to be associated with COVID-19 and we used them for further downstream analyses.

**Table 1:** Summary of TWAS and colocalization analyses in tissues and cell lines.

| Gene symbol | Tissue | Discovery: GWAS<sub>HGI</sub> |  |  |  | Validation: GWAS<sub>SCGG</sub> |  |  |  |
| --- | --- | --- | --- | --- | --- | --- | --- | --- | --- |
|  |  | TWAS<sub>z</sub> | TWAS<sub>p</sub> | PP | colocalized SNP p | TWAS<sub>z</sub> | TWAS<sub>p</sub> | PP | colocalized SNP p |
| *CXCR6* | Lung | -8.53 | 1.57×10<sup>-17</sup> | 0.79* | rs34068335<br>5.02×10<sup>-22</sup> | -4.19 | 2.84×10<sup>-5</sup> | ns | - |
|  | T follicular helper cells | - | - | 0.99** | rs35081325<br>3.82×10<sup>-39</sup> | - | - | 0.99** | rs35081325<br>2.49×10<sup>-10</sup> |
| *CCR9* | Whole blood | 6.50 | 7.90×10<sup>-11</sup> | ns | - | 6.26 | 3.78×10<sup>-10</sup> | ns | - |

GWAS<sub>HGI</sub> denotes the GWAS dataset from the Host Genetics Initiative.
GWAS<sub>SCGG</sub> represents the GWAS dataset from the Severe COVID-19 GWAS Group.
PP: posterior probability.
z: z score.
p: p-value.
\*: statistically significant by the colocalization posterior probability (CLPP) from eCAVIAR.
\*\*: statistically significant by the regional colocalization probability (RCP) from fastENLOC.
ns: no significant colocalization from either eCAVIAR or fastENLOC.
-: no available data.

**Fig 2.**
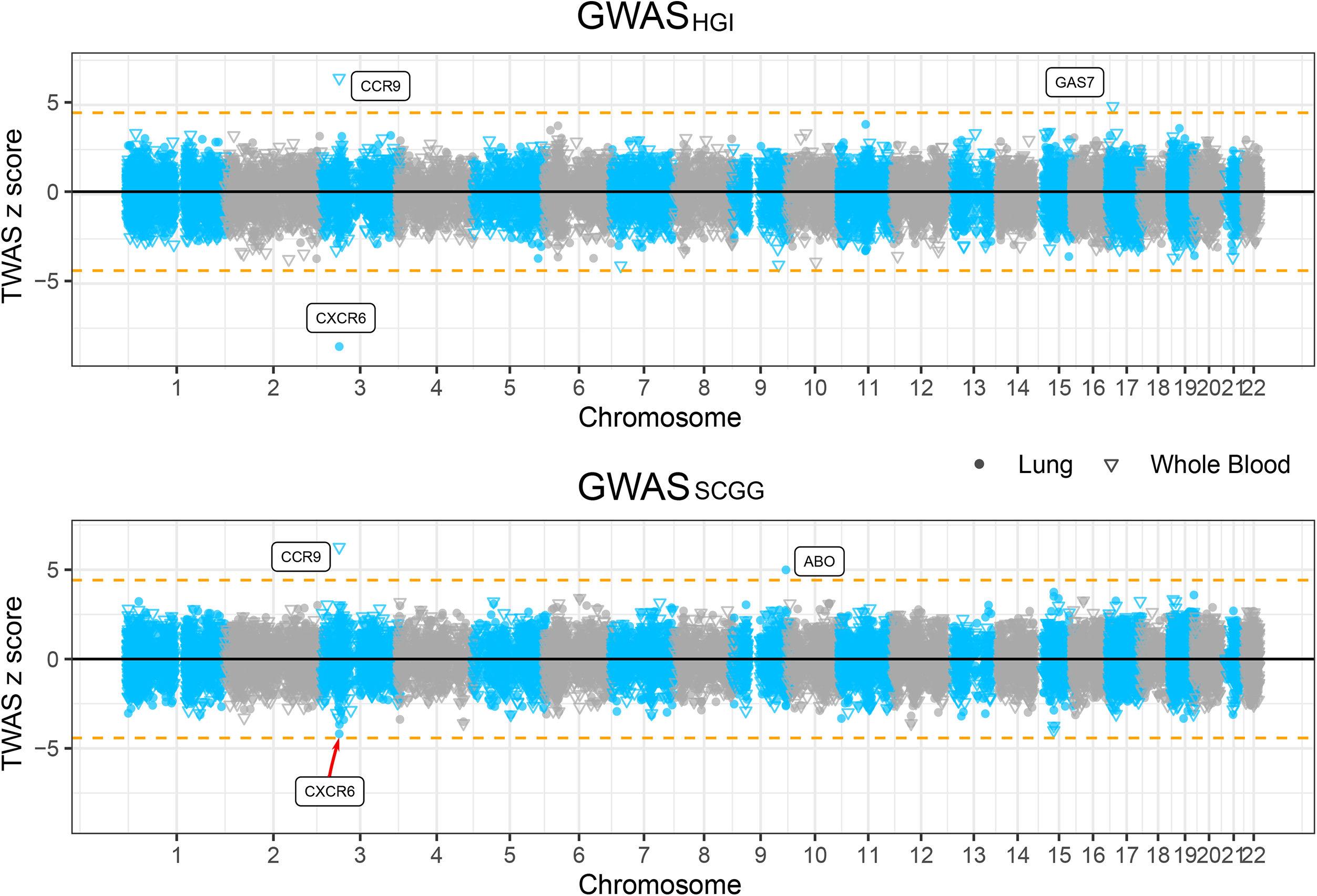
Manhattan plots illustrating the z scores of transcriptome-wide association study (TWAS) genes. TWAS z scores for two genome-wide association study (GWAS) datasets of susceptibility to severe COVID-19 using lung and whole blood tissue models. The upper panel shows the results from GWAS_HGI_ and the lower panel from GWAS_SCGG_ (see Methods). The round and triangle points denote lung and whole blood tissues, respectively, in the TWAS analysis. Dashed horizontal lines denote the Bonferroni-corrected significance threshold (|z| = 4.56, p < 5 × 10^−6^). Significant genes were highlighted with their gene symbol.

### Colocalization analysis validated the mediation effect of *CXCR6* between GWAS locus and severe COVID-19

The TWAS findings might be driven by pleiotropy or linkage effect by the LD structure in the GWAS loci instead of the true mediation effect [42] (**Fig. 3a**). To rule out the linkage effect and find further evidence of true colocalization of causal signals in the variants that were significant in both GWAS and eQTL analyses, we performed colocalization analysis by eCAVIAR and fastENLOC using several tissue-specific eQTL datasets. The eCAVIAR with the eQTL data in lung tissue revealed that the severe COVID-19 association could be mediated by the variants that were associated with the expression of *CXCR6* (CLPP = 0.79) (**Table 1**). And the colocalized SNP rs34068335 (GWAS_HGI_ p = 5.02 × 10^−22^) is also related to the increased monocyte percentage of white cells in a blood-trait GWAS study using Phenoscanner [43-45]. The fastENLOC analysis showed a high RCP between the expression of *CXCR6* in T follicular helper cells and GWAS signal in both the GWAS_HGI_ cohort (RCP=0.99) and the GWAS_SCGG_ cohort (RCP = 0.99) (**Table 1**). However, colocalization analysis of *CCR9* did not suggest strong colocalization evidence (CLPP < 0.1 and RCP < 0.1).

**Fig 3.**
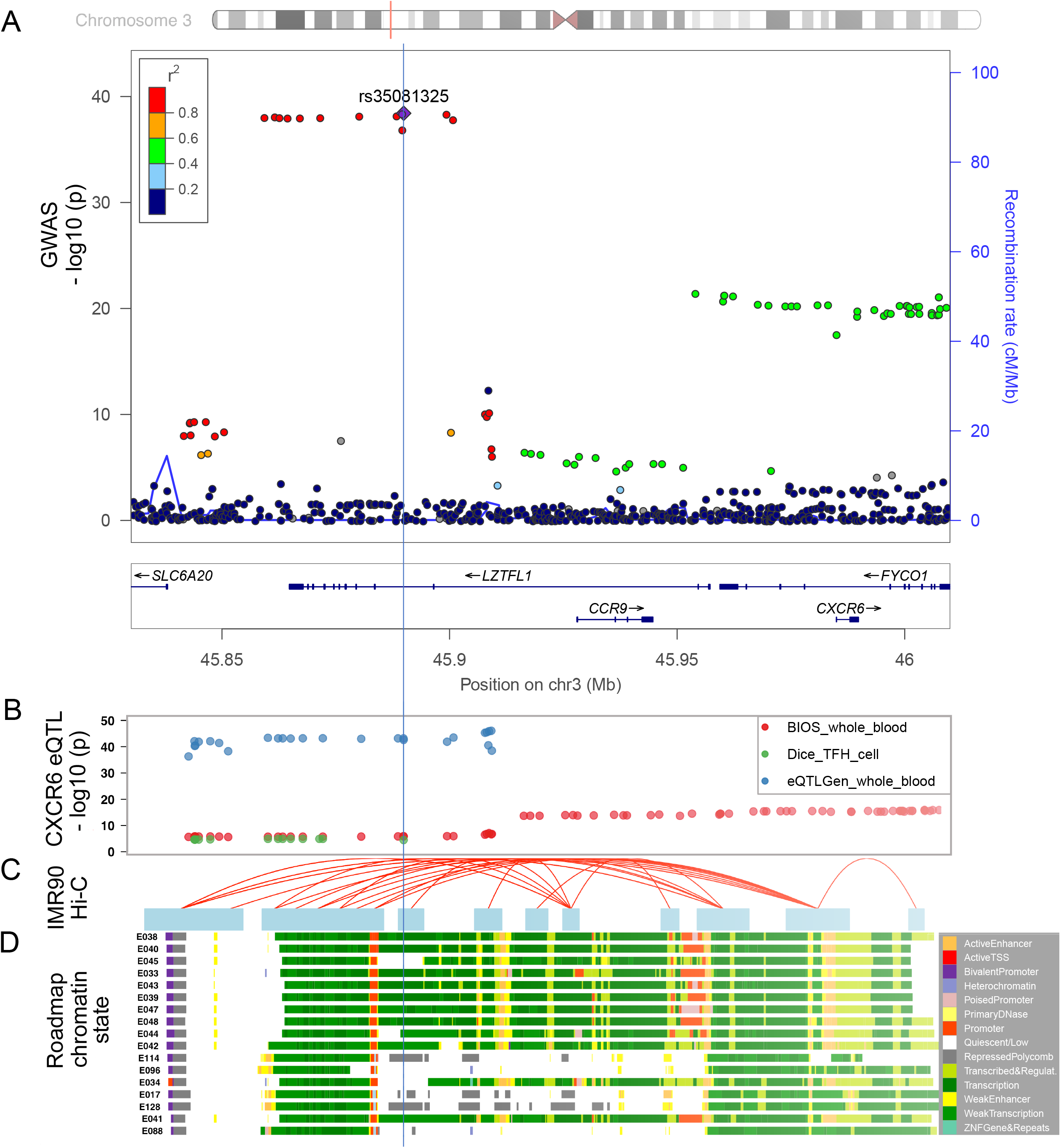
Functional genomic annotation on *3p21*.*31* locus with signals from GWAS_HGI_. (**a**) LocusZoom view of the association signals of SNPs at the *3p21*.*31* locus of GWAS_HGI_. The x-axis is the chromosome position in million base pairs (Mb) on GRCh37 reference genome and y-axis represents the –log_10_ (p-value) from GWAS_HGI_ dataset. The color indicates the strength of linkage disequilibrium from the lead SNP rs35081325. The genes within the region are annotated in the lower panel. A vertical blue line labels the position of the lead SNP rs35081325 to denote the relationship of GWAS variants to other datasets: expression quantitative trait (eQTL) (Fig. 3b), chromatin interaction (Fig. 3c), and imputed Roadmap functional elements (Fig. 3d). (**b**) The significant eQTLs associated with *CXCR6* expression in this region. The *cis-*eQTL datasets include two whole blood datasets [Biobank-based Integrative Omics Studies (BIOS) QTL and eQTLGen] and one T follicular helper cell dataset (DICE). The y axis represents the –log_10_ (p-value) from the eQTL studies. (**c**) The significant Hi-C interactions in normal lung fibroblast cell line (IMR90). Blue blocks denote the target and bait regions, and red arcs indicate the interactions between functional elements. (**d**) The region annotated with the chromatin-state segmentation track (ChromHMM) from the Roadmap Epigenomics data for T-cell and lung tissue. The Roadmap Epigenomics cell line IDs are shown on the left side: E017 (IMR90 fetal lung fibroblasts Cell Line), E033 (Primary T Cells from cord blood), E034 (Primary T Cells from blood), E038 (Primary T help naïve cells from peripheral blood), E039 (Primary T helper naïve cells from peripheral blood), E040 (Primary T helper memory cells from peripheral blood 1), E041 (Primary T helper cells PMA-Ionomycin stimulated), E042 (Primary T helper 17 cells PMA-Ionomycin stimulated), E043 (Primary T helper cells from peripheral blood), E044 (Primary T regulatory cells from peripheral blood), E045 (Primary T cells effector/memory enriched from peripheral blood), E047 (Primary T CD8 naïve cells from peripheral blood), E048 (Primary T CD8 memory cells from peripheral blood), E088 (Fetal lung), E096 (Lung), E114 (A549 EtOH 0.02pct Lung Carcinoma Cell Line), and E128 (NHLF Human Lung Fibroblast Primary Cells). The colors denote chromatin states imputed by ChromHMM, with the color key in the gray box (Methods).

### Multi-level functional annotations linked *3p21*.*31* locus with *CXCR6* and *CCR9* functions

To explore the potential functions linked with the GWAS risk variants, we examined the functional genomic annotations in this locus. Specifically, we found a consistent decreasing effect of *CXCR6* expression in T cells and whole blood from the two large-scaled eQTL datasets (**Fig. 3b**). Furthermore, multiple SNPs at the *3p21*.*31* locus reside in the annotated regulatory elements across blood, T cell, and lung cell lines (**Fig. 3c**, Methods). The Hi-C cell line data from lung fibroblast [28] also showed a significant interaction between the *3p21*.*31* locus had interactions with both *CXCR6* and *CCR9* promoter regions (**Fig. 3d**). Overall, these results from the multiple lines of evidence all supported the potential regulatory effects of the *3p21*.*31* locus on *CXCR6* expression.

### *CXCR6* differentially expressed in T_RM_ cells of severe and moderate patients

According to our tissue cell-type-specific expression database (CSEA-DB), *CXCR6* is mainly expressed in immune cells in human lung tissue (e.g., T cell and NK cell) [16]. In Liao et al.’s work, the authors reported that *CXCR6* had lower expression in severe patients than moderate patients, indicating a potential protective effect in T cells of human respiratory systems [17]. However, T cells have various resident and circulating subtypes with diverse functions [46]. To understand which subpopulation(s) of T cells might be associated with the severity of COVID-19, we used the BLAF scRNA-seq data of six severe patients and three moderate patients. The data included 6,491 T-cells (4,356 from six severe patients and 2,135 from three moderate patients). We further used a set of 31 T_RM_ cell marker genes to distinguish the T_RM_ cells and conventional CD8^+^ T cells (Methods). As shown in **Fig. 4a and 4b**, the T_RM_ cells and conventional T cells could be distinguished in both moderate and severe patients with the classic T_RM_ cells markers (*CXCR6* [31], CD69 [47], *ITGAE* (the gene encoding CD103) [47, 48], *ZNF683* [48], and *XCL1* [46]) and three negative-control markers (*SELL* (the gene encoding CD62L) [47], KLF2, and S1PR1 [49]) from previous study [31]. Among the 1,090 lung T_RM_ cells, we found that 675 cells were from moderate patients and only 415 cells were from severe patients. This represented a 3.32-fold decrease for the expected number of T_RM_ cells in severe patients. We used the non-parametric Wilcoxon rank sum test to identify the DEGs in the T_RM_ cells between severe and moderate patients and found *CXCR6* had significantly lower expression in the severe patients than the moderate patients (p < 2.5 × 10^−16^, fold change = 1.57, **Fig. 4c**).

**Fig 4.**
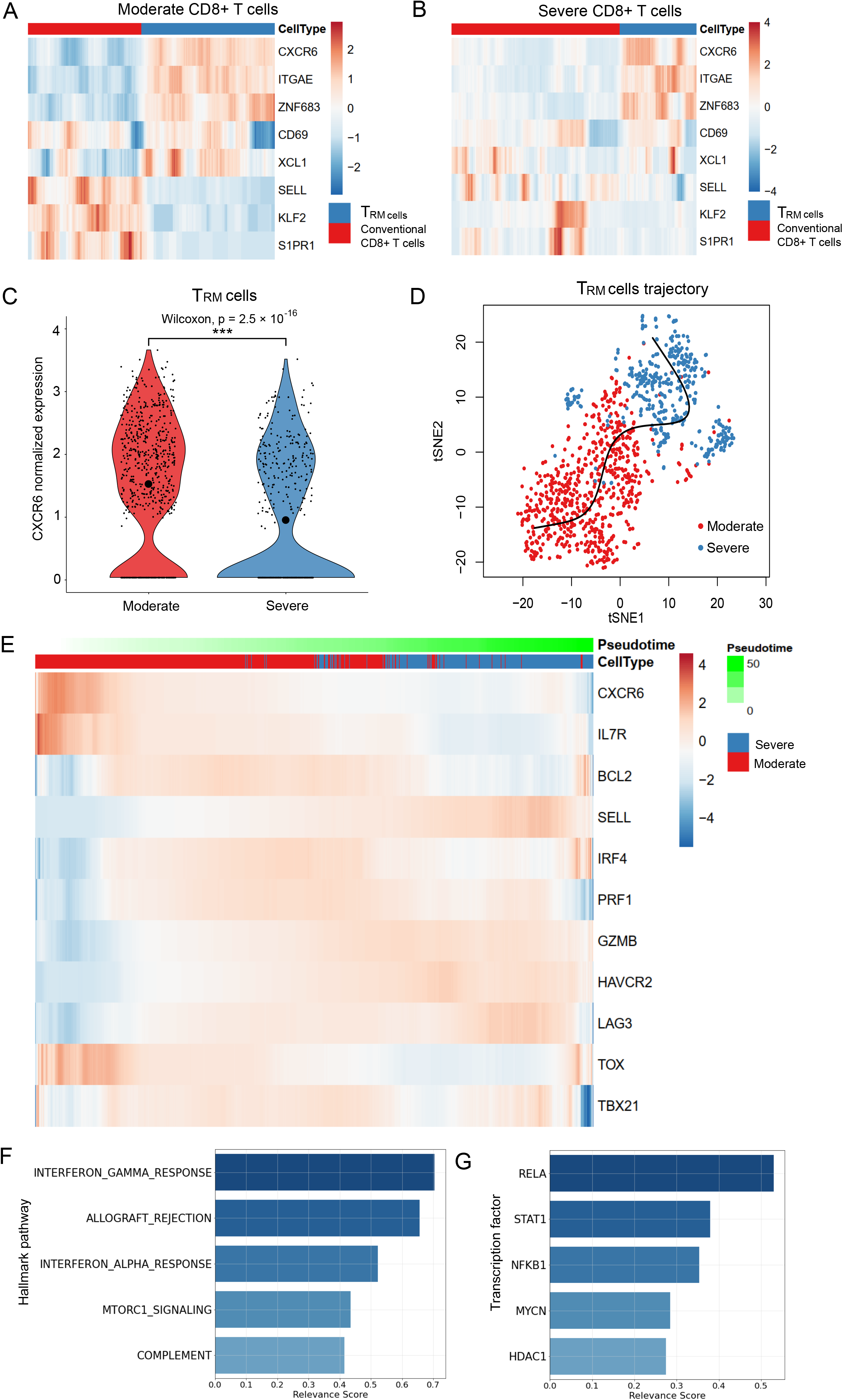
Single cell transcriptome analysis of the severe and moderate COVID-19 patients. (**a**) Relative expression of the lung resident memory CD8^+^ T (T_RM_) signature genes in T_RM_ cells and conventional CD8^+^ T cells in moderate patients. (**b**) Relative expression of the T_RM_ featured genes in T_RM_ cells and conventional CD8^+^ T cells in severe patients. (**c**) *CXCR6* expression in the T_RM_ cells of moderate and severe patients. We split the T_RM_ cells from the annotation of the original paper with 31 marker genes (Methods). We conducted a two-sided non-parameter Wilcoxon rank sum test to test whether *CXCR6* was differentially expressed in moderate (red) and severe (blue) groups of T_RM_ cells. “***” indicates it is genome-wide significant after multiple-test correction of all expressed genes. The small points denote the normalized expression in each cell. Mean normalized expression of *CXCR6* in each group is highlighted with the largest circle in black. (**d**) Pseudotime inference for the moderate and severe T_RM_ cells. The red and blue points on t-Distributed Stochastic Neighbor Embedding (tSNE) projection denote the T_RM_ cells from moderate and severe patients, respectively. The x-axis and y-axis are the first and second dimension of the tSNE, respectively. (**e**) Relative expression of the *CXCR6* and naïve and effector T cell markers along the pseudotime proportional to the green color. The gene expressions are scaled by cells. Cells from moderate and severe groups are annotated in blue and red. (**f**) Relevance score for hallmark pathways from the molecular signatures database (MSigDB) along the pseudotime. The relevance score (R^2^ coefficient of determination) indicates the proportion of variance in the pseudotime explained by the genes in the hallmark pathways. (**g**) Relevance score for transcription factors and their target genes along the pseudotime. The relevance score denotes the proportion of variance in the pseudotime explained by the target genes regulated by the transcription factor.

### Inferring the transcriptional programs that drive the cell status transition

To understand the transition between moderate and severe T_RM_ cells, we constructed the cell trajectory/pseudotime along with T_RM_ cells by using Slingshot (**Fig. 4d**) [34]. Next, we applied our DrivAER approach (Driving transcriptional programs based on AutoEncoder derived Relevance scores) [36] to identify the potential transcriptional programs that were most likely involved in the cell trajectory/pseudotime. **Fig. 4e** shows a scaled heatmap to demonstrate the relative expression of naïve and effector markers of T cells in the order of pseudotime generated by Slingshot [34, 39]. We identified that the severe T_RM_ cells were mainly gathered in the later stage of the pseudotime. The naïve markers (*IL7R, BCL2*) were higher expressed in moderate patients than in severe patients (except *SELL*). On the contrary, some effector markers (*GZMB, HAVCR2, LAG3, IFNG*) were lower expressed in moderate patients than in severe patients. Other effector markers (*IRF4, PRF1*) had higher expression in the middle of the transition than their expression at the start and end sides. These results indicated that the T_RM_ cells in severe patients still in pro-inflammatory status although the T_RM_ cells status were more heterogeneous in severe patients than in moderate patients (**Fig. 4a, 4b, and 4e**). As shown in **Fig. 4f and 4g**, the top five molecular signatures (relevance score > 0.25) identified by DrivAER included T-cell pro-inflammatory actions (interferon gamma response, allograft rejection [50], interferon alpha response, and complement system) as well as proliferative mTORC1 signaling pathway [51]. Among the top TFs (relevance score > 0.25) that drove this cell trajectory, the DNA binding RELA-NFKB1 complex is involved in several biological processes, such as inflammation, immunity, and cell growth initiated by external stimuli. The signal transducer and activator of transcription (*STAT1*) and its regulator histone deacetylase (*HDAC1*) could be activated by various ligands including interferon-alpha and interferon-gamma. In summary, the TF results are well consistent with our previous hallmark pathway findings (**Additional file: Table S1 and Table S2**).

### Several genome-wide significant SNPs might change the TF binding site affinity

To understand the potential TF binding affinity changes of genome-wide significant SNPs, we conducted the DNA motif recognition analysis of the seven TFs related to the transcriptional program between moderate and severe T_RM_ cells (relevance score > 0.25, **Additional file 1: Table S2**). We identified SNP rs10490770 [T/C, minor allele frequency (MAF) = 0.097, GWAS_HGI_ = 9.53 × 10^−39^] and SNP rs67959919 (G/A, MAF = 0.097, GWAS_HGI_ = 8.83 × 10^−39^) that were predicted to alter the binding affinity of TFs RELA and SP1, respectively (**Additional file 1: Fig. S1a and S1b**). Moreover, these two SNPs were in the high LD region (r^2^ > 0.8) with several significant lead eQTLs (SNP rs35896106 and rs17713054) of *CXCR6* in whole blood (p = 5.03 × 10^−37^) and T follicular helper cell (p = 1.30 × 10^−5^) (**Fig. 3b**). In summary, the genome-wide significant SNPs were predicted to change the binding affinity of those TFs highly related to T_RM_ cells status transition, (**Additional file 2: Table S3**), suggesting their potential regulation of *CXCR6* expression.

## Discussion

In this work, we developed a multi-level, integrative genetic and functional analysis framework to explore the host genetic factors on the expression change of GWAS-implicated genes for COVID-19 severity. Specifically, we conducted TWAS analysis for two independent COVID-19 GWAS datasets. We identified and replicated two chemokine receptor genes, *CXCR6* and *CCR9*, with a protective effect in the lung and a risk effect in whole blood, respectively. *CXCR6* is expressed in T lymphocytes and essential genes in CD8^+^ T_RM_ cells, mediating the homing of T_RM_ cells to the lung along with its ligand *CXCL16* [52, 53]. *CCR9* was reported to regulate chemotaxis in response to thymus-expressed chemokine in T cells [54]. The colocalization analysis identified that both GWAS and eQTLs of *CXCR6* had high colocalization probabilities in the lung, whole blood, and T follicular helper cells, which confirms the genetic regulation roles at this locus. At the single cell level, our DEG analysis identified *CXCR6* gene had lower expression in the COVID-19 severe patients than the moderate patients in both T cells and T_RM_ cells, supporting its protective effect identified in TWAS analysis in lung and whole blood. The expected proportion of T_RM_ cells also decreased by 3.32-fold (**Table 2**). Interestingly, these findings were replicated in circulating CXCR6^+^ CD8^+^ T cells of severe and control/mild patients by flow cytometry experiment [53]. We identified the major transition force from moderate T_RM_ cells to severe T_RM_ cells are pro-inflammatory pathways and TFs.

**Table 2:** Counts and ratio of T_RM_ cells in moderate and severe patients.

| Patient group<br>(sample size) | # CD8<sup>+</sup> T<br>cells | # T<sub>RM</sub><br>cells | T<sub>RM</sub> cell proportion ratio<br>(Moderate/Severe) |
| --- | --- | --- | --- |
| Moderate (3) | 2,135 | 675 | 3.32 |
| Severe (6) | 4,356 | 415 |  |

#: the counted number.

From the TWAS and colocalization analysis in lung and immune cells, we successfully replicated that *CXCR6* was centered in the GWAS signal at locus *3p21*.*31*. Previous studies have reported that CXCR6^-/-^ significantly decreases airway lung T_RM_ cells due to altered trafficking of CXCR6^-/-^ cells within the lung of the mice [52], which could explain a much less proportion of T_RM_ cells in severe patients than moderate patients. The lung T_RM_ cells provide the first line of defense against infection and coordinate the subsequent adaptive response [55]. The previous study has reported that T_RM_ cells constitutively expressed surface receptors (PD-1 and CTLA-4) that are associated with inhibition of T cell function, which might prevent excessive activation or inflammation in the tissue niche [56].

We further used nine classic naïve markers (e.g., *BCL2, SELL, TCF7*, and *IL7R)* and ten classic effector markers (e.g., *GZMB, PRF1, IFNG, LAG3*, and *PDCD1*) to quantify the naïve and effector status of the T_RM_ cells (**Additional file 1: Fig. S2**). T_RM_ cells in severe patients had a much higher median of effector marker score (0.44 in severe and 0.18 in moderate T_RM_ cells) than T_RM_ cells in moderate patients did, suggesting that the severe T_RM_ cells had much higher activities in inflammation as we discovered in **Fig. 4f** despite their proportion decrease. For the naïve score (**Additional file 1: Fig. S2**), both moderate and severe T_RM_ cells had limited expressions (median score: 0.028 in severe and 0.038 in moderate T_RM_ cells). Interestingly, if we removed the lymph node homing receptor *SELL* [31] from the naïve markers list, we would find the median score in severe naïve markers would drop to 0 (**Additional file 1: Fig. S2**). This indicated that *SELL* expression contributed greatly to the naïve status of T_RM_ severe patients. Consistently in **Fig. 4e**, we could also observe that a large proportion of T_RM_ cells had higher *SELL* expression in severe patients than in moderate patients, suggesting the T_RM_ cells in severe patients might not be in a stable cell status due to the lymph node homing signal (*SELL*). To this end, we hypothesized that genetically lower expressed *CXCR6* would decrease the proportion of T_RM_ cells residing in the lung through the CXCR6/CXCL16 axis [52, 53], impairing the first-line defense. Moreover, the lower expression of *CXCR6* would also lead to the “unstable” residency of T_RM_ cells in lung (**Fig. 4b**). The T_RM_ cells play essential roles for orchestrating the immune system, lack of which would lead to severe COVID-19 symptoms, such as acute respiratory distress syndrome, cytokine storm and major multi-organ damage [57] (**Fig. 5**).

**Fig 5.**
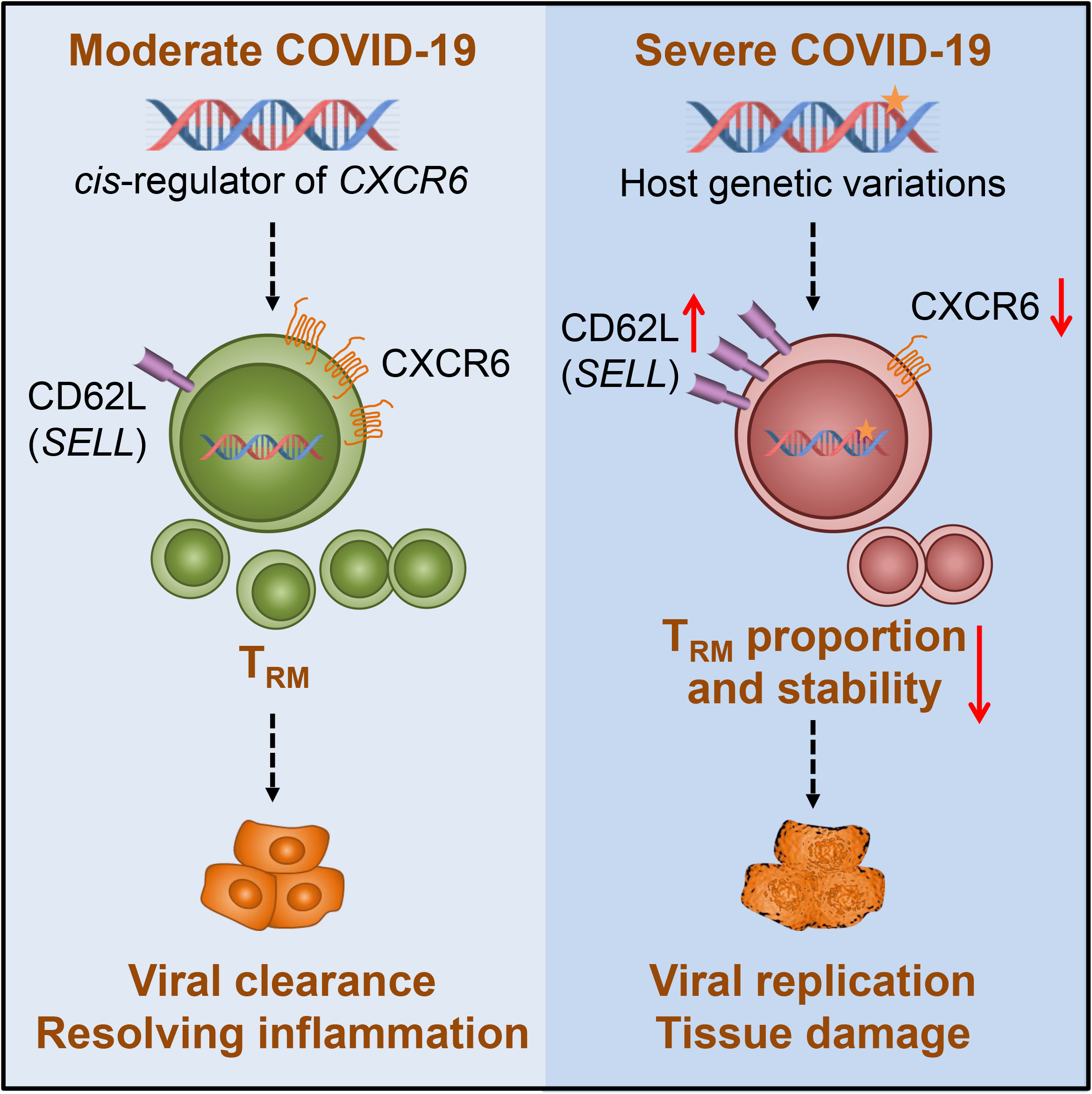
The proposed *CXCR6* regulation mechanism on COVID-19 severity. We proposed one pathogenesis mechanism using current knowledge to explain how the lower expression of *CXCR6* could be associated with the outcome of severe COVID-19 symptoms, which was supported by our findings of the genetic factors on decreasing the *CXCR6* expression and aligned with our observations from single cell transcriptome analysis. The star on the DNA indicates the host genetic effects.

In this study, we mainly focused on the multi-evidence validated gene *CXCR6* and its mechanism related to severe COVID-19. Although we are unable to directly test the genotype of those severe patients, the association of the single cell level phenotype (lower expression of CXCR6 and decreased proportion of CD8^+^ CXCR6^+^ T cells) and the severe COVID-19 has been observed in another work with flow cytometry experiments [53]. We are aware of the genetic factors on *CXCR6* might only explain a proportion of the severe COVID-19 variance. Other genetic mechanisms discovered in GWAS and TWAS analyses need further exploration [6]. The GWAS_HGI_ dataset used in this study was HGI round 4 (alpha), which was the largest GWAS by the access date of October 20, 2020. However, it was not the currently largest GWAS meta-analysis for severe COVID-19 when we prepared the manuscript. This research field is evolving very fast, due to the urgent demand of public health. Currently, the largest GWAS HGI round 4 (freeze) contained more samples (4,336 cases/ 353,891 controls), and it included two independent datasets we used in this study. Considering that the GWAS_HGI_ dataset included ∼10% control samples from the Asian population, we checked the LocusZoom plot of the chr3: 45.80-46.40 million base pairs (Mb) region on GRCh37 reference genome. We found a consistent tendency in GWAS round 4 alpha and freeze version (**Additional file1: Fig. S3**). Another limitation is that the scRNA-seq data only had nine COVID-19 patient samples (six severe and three moderate samples), which might not provide enough statistical power at the sample level as it is commonly considered each scRNA-seq data acts like a population. Finally, the TF binding site affinity alterations were assessed based on computational prediction, therefore, the *in vivo* effects require experimental validation. We anticipate more and larger datasets will be released in the near future. We will apply our integrative analysis approach to such new data.

## Conclusions

Our work systematically explored the genetic effect on gene expression at chromosome locus *3p21*.*31* and pinpointed the gene *CXCR6* might be involved in the severity of COVID-19. Several genome-wide significant SNPs were within the LD block of *CXCR6* eQTLs in immune-related cells. In a scRNA-seq COVID-19 BALF dataset, we characterized that *CXCR6* (T_RM_ cells marker gene) had a lower expression in severe patients than in moderate patients. Moreover, the T_RM_ cells in severe patients had a 3.32-fold proportion decrease and much higher pro-inflammatory activity than T_RM_ cells in moderate patients. Based on these observations, we proposed a potential mechanism on how the lower expression of *CXCR6* regulated by the endogenous factors could progress to severe COVID-19 outcomes.

## List of abbreviations

BALF: bronchoalveolar lavage fluid
BIOS: Biobank-based Integrative Omics Studies
ChromHMM: chromatin-state hidden Markov model
COVID-19: coronavirus disease 2019
CLPP: colocalization posterior probability
CSEA-DB: cell-type-specific expression database
DAP: deterministic approximation of posteriors
DEG: differentially expressed gene
DICE: database of immune cell expression
DrivAER: Driving transcriptional programs based on AutoEncoder derived Relevance scores
eQTL: expression quantitative trait
GReX: genetically regulated expression
GWAS: genome-wide association study
HGI: Host Genetics Initiative
Hi-C: high-throughput chromatin interaction
LD: linkage disequilibrium
MAF: minor allele frequency
MASHR: multivariate adaptive shrinkage in R
Mb: million base pairs
MSigDB: molecular signatures database
PIP: posterior inclusion probability
PWM: position weight matrix
SARS-CoV-2: severe acute respiratory syndrome coronavirus 2
RCP: regional colocalization probability
SCGG: Severe COVID-19 GWAS Group
scRNA-seq: single cell RNA sequencing
tSNE: t-Distributed Stochastic Neighbor Embedding
TF: transcription factor
T_RM_cells: resident memory CD8+ T cells
TWAS: transcriptome-wide association study

## Acknowledgements

We appreciate Drs. Teng Liu and Dawei Zou for the valuable comments. We thank all members of the Bioinformatics and Systems Medicine Laboratory for the discussion.

## Additional files

Additional file 1.pdf: Fig S1: Sequence logos representing DNA binding site generated from position weight matrix (PWM) for transcription factor RELA and SP1. Fig. S2. Violin plots showing the distribution of key features between moderate and severe patients. Fig. S3. LocusZoom views for two Host Genetics Initiates GWAS datasets at *3p21*.*31* locus. Table S1: Hallmark pathways and their relevance scores. Table S2: Transcription factors and their relevance scores.

Additional file 2.xls: Table S3: Predicted transcription factors (SP1 and RELA) bind affinity alterations on genome-wide significant SNPs at locus *3p21*.*31*.

## References

1. Zhao Z, Li H, Wu X, Zhong Y, Zhang K, Zhang YP, Boerwinkle E, Fu YX: Moderate mutation rate in the SARS coronavirus genome and its implications. BMC Evol Biol 2004, 4:21.

2. Liu S, Shen J, Fang S, Li K, Liu J, Yang L, Hu CD, Wan J: Genetic Spectrum and Distinct Evolution Patterns of SARS-CoV-2. Front Microbiol 2020, 11:593548.

3. Wu Z, McGoogan JM: Characteristics of and Important Lessons From the Coronavirus Disease 2019 (COVID-19) Outbreak in China: Summary of a Report of 72314 Cases From the Chinese Center for Disease Control and Prevention. JAMA 2020, 323:1239–1242.

4. Bhopal SS, Bhopal R: Sex differential in COVID-19 mortality varies markedly by age. vol. 396. pp. 532–533: Lancet Publishing Group; 2020:532-533.

5. Severe Covid GG, Ellinghaus D, Degenhardt F, Bujanda L, Buti M, Albillos A, Invernizzi P, Fernandez J, Prati D, Baselli G, et al: Genomewide Association Study of Severe Covid-19 with Respiratory Failure. N Engl J Med 2020, 383:1522–1534.

6. Pairo-Castineira E, Clohisey S, Klaric L, Bretherick AD, Rawlik K, Pasko D, Walker S, Parkinson N, Fourman MH, Russell CD, et al: Genetic mechanisms of critical illness in Covid-19. Nature 2020.

7. GTEx Consortium: The GTEx Consortium atlas of genetic regulatory effects across human tissues. Science 2020, 369:1318–1330.

8. Võsa U, Claringbould A, Westra H-J, Bonder MJ, Deelen P, Zeng B, Kirsten H, Saha A, Kreuzhuber R, Kasela S, et al: Unraveling the polygenic architecture of complex traits using blood eQTL metaanalysis. bioRxiv 2018:447367.

9. Schmiedel BJ, Singh D, Madrigal A, Valdovino-Gonzalez AG, White BM, Zapardiel-Gonzalo J, Ha B, Altay G, Greenbaum JA, McVicker G, et al: Impact of Genetic Polymorphisms on Human Immune Cell Gene Expression. Cell 2018, 175:1701–1715 e1716.

10. Gamazon ER, Wheeler HE, Shah KP, Mozaffari SV, Aquino-Michaels K, Carroll RJ, Eyler AE, Denny JC, Consortium GT, Nicolae DL, et al: A gene-based association method for mapping traits using reference transcriptome data. Nat Genet 2015, 47:1091–1098.

11. Giambartolomei C, Vukcevic D, Schadt EE, Franke L, Hingorani AD, Wallace C, Plagnol V: Bayesian test for colocalisation between pairs of genetic association studies using summary statistics. PLoS Genet 2014, 10:e1004383.

12. Dai Y, Pei G, Zhao Z, Jia P: A Convergent Study of Genetic Variants Associated With Crohn’s Disease: Evidence From GWAS, Gene Expression, Methylation, eQTL and TWAS. Front Genet 2019, 10:318.

13. Dai Y, Hu R, Pei G, Zhang H, Zhao Z, Jia P: Diverse types of genomic evidence converge on alcohol use disorder risk genes. J Med Genet 2020, 57:733–743.

14. Mathys H, Davila-Velderrain J, Peng Z, Gao F, Mohammadi S, Young JZ, Menon M, He L, Abdurrob F, Jiang X, et al: Single-cell transcriptomic analysis of Alzheimer’s disease. Nature 2019, 570:332–337.

15. Papalexi E, Satija R: Single-cell RNA sequencing to explore immune cell heterogeneity. Nat Rev Immunol 2018, 18:35–45.

16. Dai Y, Hu R, Manuel AM, Liu A, Jia P, Zhao Z: CSEA-DB: an omnibus for human complex trait and cell type associations. Nucleic Acids Res 2021, 49:D862–D870.

17. Liao M, Liu Y, Yuan J, Wen Y, Xu G, Zhao J, Cheng L, Li J, Wang X, Wang F, et al: Single-cell landscape of bronchoalveolar immune cells in patients with COVID-19. Nat Med 2020, 26:842–844.

18. Barbeira AN, Dickinson SP, Bonazzola R, Zheng J, Wheeler HE, Torres JM, Torstenson ES, Shah KP, Garcia T, Edwards TL, et al: Exploring the phenotypic consequences of tissue specific gene expression variation inferred from GWAS summary statistics. Nature Communications 2018, 9:1–20.

19. Urbut SM, Wang G, Carbonetto P, Stephens M: Flexible statistical methods for estimating and testing effects in genomic studies with multiple conditions. Nat Genet 2019, 51:187–195.

20. Hormozdiari F, van de Bunt M, Segrè AV, Li X, Joo JWJ, Bilow M, Sul JH, Sankararaman S, Pasaniuc B, Eskin E: Colocalization of GWAS and eQTL Signals Detects Target Genes. American Journal of Human Genetics 2016, 99:1245–1260.

21. Wen X, Pique-Regi R, Luca F: Integrating molecular QTL data into genome-wide genetic association analysis: Probabilistic assessment of enrichment and colocalization. PLOS Genetics 2017, 13:e1006646–e1006646.

22. Genomes Project C, Auton A, Brooks LD, Durbin RM, Garrison EP, Kang HM, Korbel JO, Marchini JL, McCarthy S, McVean GA, Abecasis GR: A global reference for human genetic variation. Nature 2015, 526:68–74.

23. Lee Y, Luca F, Pique-Regi R, Wen X: Bayesian Multi-SNP genetic association analysis: Control of FDR and use of summary statistics. pp. 316471–316471: bioRxiv; 2018:316471-316471.

24. Võsa U, Claringbould A, Westra HJ, Bonder MJ, Deelen P, Zeng B, Kirsten H, Saha A, Kreuzhuber R, Kasela S, et al: Unraveling the polygenic architecture of complex traits using blood eQTL meta-analysis. vol. 18. pp. 10–10: bioRxiv; 2018:10-10.

25. Zhernakova DV, Deelen P, Vermaat M, van Iterson M, van Galen M, Arindrarto W, van ’t Hof P, Mei H, van Dijk F, Westra HJ, et al: Identification of context-dependent expression quantitative trait loci in whole blood. Nat Genet 2017, 49:139–145.

26. Ernst J, Kellis M: ChromHMM: automating chromatin-state discovery and characterization. Nat Methods 2012, 9:215–216.

27. Roadmap Epigenomics C, Kundaje A, Meuleman W, Ernst J, Bilenky M, Yen A, Heravi-Moussavi A, Kheradpour P, Zhang Z, Wang J, et al: Integrative analysis of 111 reference human epigenomes. Nature 2015, 518:317–329.

28. Dixon JR, Selvaraj S, Yue F, Kim A, Li Y, Shen Y, Hu M, Liu JS, Ren B: Topological domains in mammalian genomes identified by analysis of chromatin interactions. Nature 2012, 485:376–380.

29. Hahne F, Ivanek R: Visualizing genomic data using Gviz and bioconductor. In Volume 1418: Humana Press Inc.; 2016: 335–351

30. Liu T, Jia P, Fang B, Zhao Z: Differential Expression of Viral Transcripts From Single-Cell RNA Sequencing of Moderate and Severe COVID-19 Patients and Its Implications for Case Severity. Front Microbiol 2020, 11:603509.

31. Kumar BV, Ma W, Miron M, Granot T, Guyer RS, Carpenter DJ, Senda T, Sun X, Ho SH, Lerner H, et al: Human Tissue-Resident Memory T Cells Are Defined by Core Transcriptional and Functional Signatures in Lymphoid and Mucosal Sites. Cell Rep 2017, 20:2921–2934.

32. Pont F, Tosolini M, Fournie JJ: Single-Cell Signature Explorer for comprehensive visualization of single cell signatures across scRNA-seq datasets. Nucleic Acids Res 2019, 47:e133.

33. Stuart T, Butler A, Hoffman P, Hafemeister C, Papalexi E, Mauck WM, 3rd, Hao Y, Stoeckius M, Smibert P, Satija R: Comprehensive Integration of Single-Cell Data. Cell 2019, 177:1888–1902 e1821.

34. Street K, Risso D, Fletcher RB, Das D, Ngai J, Yosef N, Purdom E, Dudoit S: Slingshot: cell lineage and pseudotime inference for single-cell transcriptomics. BMC Genomics 2018, 19:477.

35. Hastie T, Stuetzle W: Principal Curves. Journal of the American Statistical Association 1989, 84:502–516.

36. Simon LM, Yan F, Zhao Z: DrivAER: Identification of driving transcriptional programs in single-cell RNA sequencing data. Gigascience 2020, 9.

37. Eraslan G, Simon LM, Mircea M, Mueller NS, Theis FJ: Single-cell RNA-seq denoising using a deep count autoencoder. Nat Commun 2019, 10:390.

38. Liberzon A, Birger C, Thorvaldsdottir H, Ghandi M, Mesirov JP, Tamayo P: The Molecular Signatures Database (MSigDB) hallmark gene set collection. Cell Syst 2015, 1:417–425.

39. Han H, Cho JW, Lee S, Yun A, Kim H, Bae D, Yang S, Kim CY, Lee M, Kim E, et al: TRRUST v2: an expanded reference database of human and mouse transcriptional regulatory interactions. Nucleic Acids Res 2018, 46:D380–D386.

40. Nguyen NTT, Contreras-Moreira B, Castro-Mondragon JA, Santana-Garcia W, Ossio R, Robles-Espinoza CD, Bahin M, Collombet S, Vincens P, Thieffry D, et al: RSAT 2018: regulatory sequence analysis tools 20th anniversary. Nucleic Acids Res 2018, 46:W209–W214.

41. Weirauch MT, Yang A, Albu M, Cote AG, Montenegro-Montero A, Drewe P, Najafabadi HS, Lambert SA, Mann I, Cook K, et al: Determination and inference of eukaryotic transcription factor sequence specificity. Cell 2014, 158:1431–1443.

42. Wainberg M, Sinnott-Armstrong N, Mancuso N, Barbeira AN, Knowles DA, Golan D, Ermel R, Ruusalepp A, Quertermous T, Hao K, et al: Opportunities and challenges for transcriptome-wide association studies. Nat Genet 2019, 51:592–599.

43. Astle WJ, Elding H, Jiang T, Allen D, Ruklisa D, Mann AL, Mead D, Bouman H, Riveros-Mckay F, Kostadima MA, et al: The Allelic Landscape of Human Blood Cell Trait Variation and Links to Common Complex Disease. Cell 2016, 167:1415-1429.e1419.

44. Kamat MA, Blackshaw JA, Young R, Surendran P, Burgess S, Danesh J, Butterworth AS, Staley JR: PhenoScanner V2: An expanded tool for searching human genotype-phenotype associations. Bioinformatics 2019, 35:4851–4853.

45. Staley JR, Blackshaw J, Kamat MA, Ellis S, Surendran P, Sun BB, Paul DS, Freitag D, Burgess S, Danesh J, et al: PhenoScanner: A database of human genotype-phenotype associations. Bioinformatics 2016, 32:3207–3209.

46. Hombrink P, Helbig C, Backer RA, Piet B, Oja AE, Stark R, Brasser G, Jongejan A, Jonkers RE, Nota B, et al: Programs for the persistence, vigilance and control of human CD8(+) lung-resident memory T cells. Nat Immunol 2016, 17:1467–1478.

47. Martin MD, Badovinac VP: Defining Memory CD8 T Cell. Front Immunol 2018, 9:2692.

48. Wauters E, Van Mol P, Garg AD, Jansen S, Van Herck Y, Vanderbeke L, Bassez A, Boeckx B, Malengier-Devlies B, Timmerman A, et al: Discriminating mild from critical COVID-19 by innate and adaptive immune single-cell profiling of bronchoalveolar lavages. Cell Res 2021.

49. Skon CN, Lee JY, Anderson KG, Masopust D, Hogquist KA, Jameson SC: Transcriptional downregulation of S1pr1 is required for the establishment of resident memory CD8+ T cells. Nat Immunol 2013, 14:1285–1293.

50. Benichou G, Gonzalez B, Marino J, Ayasoufi K, Valujskikh A: Role of Memory T Cells in Allograft Rejection and Tolerance. Front Immunol 2017, 8:170.

51. Yu JS, Cui W: Proliferation, survival and metabolism: the role of PI3K/AKT/mTOR signalling in pluripotency and cell fate determination. Development 2016, 143:3050–3060.

52. Wein AN, McMaster SR, Takamura S, Dunbar PR, Cartwright EK, Hayward SL, McManus DT, Shimaoka T, Ueha S, Tsukui T, et al: CXCR6 regulates localization of tissue-resident memory CD8 T cells to the airways. J Exp Med 2019, 216:2748–2762.

53. Payne DJ, Dalal S, Leach R, Parker R, Griffin S, McKimmie CS, Cook GP, Richards SJ, Hillmen P, Munir T, et al: The CXCR6/CXCL16 axis links inflamm-aging to disease severity in COVID-19 patients. bioRxiv 2021:2021.2001.2025.428125.

54. Lee HS, Kim HR, Lee EH, Jang MH, Kim SB, Park JW, Seoh JY, Jung YJ: Characterization of CCR9 expression and thymus-expressed chemokine responsiveness of the murine thymus, spleen and mesenteric lymph node. Immunobiology 2012, 217:402–411.

55. Ardain A, Marakalala MJ, Leslie A: Tissue-resident innate immunity in the lung. Immunology 2020, 159:245–256.

56. Szabo PA, Miron M, Farber DL: Location, location, location: Tissue resident memory T cells in mice and humans. Sci Immunol 2019, 4.

57. Tay MZ, Poh CM, Renia L, MacAry PA, Ng LFP: The trinity of COVID-19: immunity, inflammation and intervention. Nat Rev Immunol 2020, 20:363–374.

