## Additional file 1 for "Association of *CXCR6* with COVID-19 severity: Delineating the host genetic factors in transcriptomic regulation"

Dai et al.

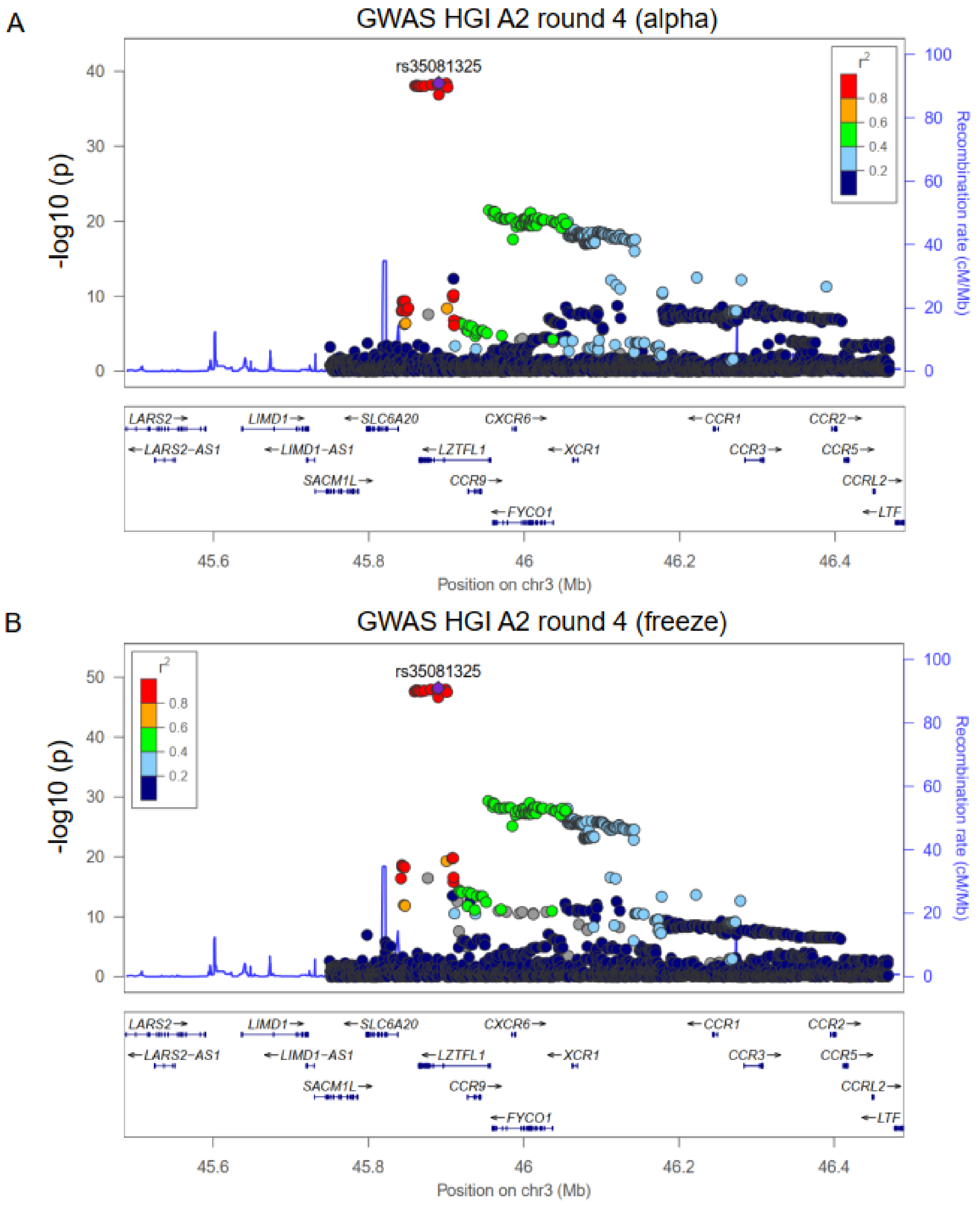


**Fig. S1. Sequence logos representing DNA binding site generated from position weight matrix (PWM) for transcription factor RELA and SP1**. In (**a**) and (**b**)**,** the SNPs rs10490770

(T/C) and rs67959919 (G/A) were predicted to have the strongest impact on their sequence (GTGGATTTTCA - Reverse strand, p = 9.8 × $10^{-4}$ and TACCCGCCGG - Reverse strand, p = 9.3 × $10^{-4}$) by utilizing the RSAT online tool (**Additional file 1: Table S1**). The polymorphism site within the transcription factor binding site is highlighted in the red box.

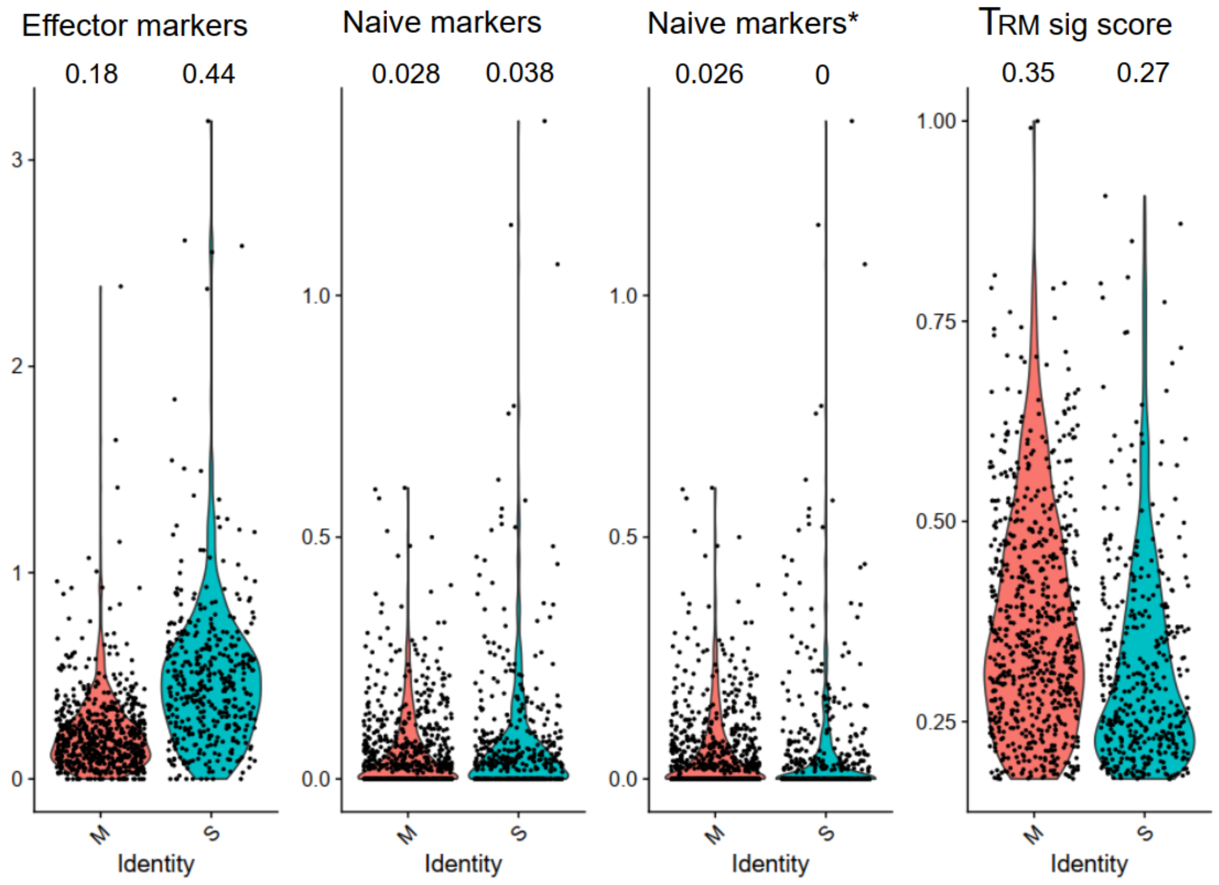


**Fig. S2. Violin plots showing the distribution of key features between moderate and severe patients.** We calculated the proportion of the effector and naïve T cell markers expressed in each cell. The naïve markers include *BCL2, SELL, KLF2, CCR7, TCF7, LEF1, ID3, BACH2*, and *IL7R*. And the effector markers include *GZMB, PRF1, IRF4, IFNG, TNFRSF9, PDCD1, LAG3, HAVCR2, TOX*, and *NR4A2*. Median score of each category is on the top of each violin plot accordingly. The "*" on the third column denotes the naïve markers without *SELL* gene. The

“$T_{RM}$ sig score” was calculated from the 31 $T_{RM}$ signature genes described in Methods. The M and S represent moderate and severe patients, respectively.

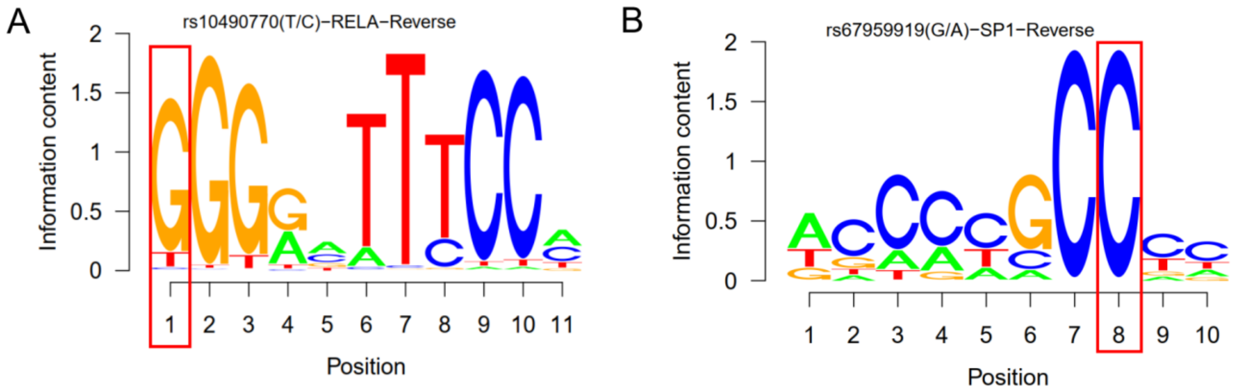


**Fig. S3. LocusZoom views for two Host Genetics Initiates GWAS datasets at *3p21.31* locus**. In (**a**) and (**b**)**,** the x-axis is the chromosome position in million base pairs (Mb) on GRCh37 reference genome and y-axis is the SNP $-\log_{10}$ (p-value) from the two $GWAS_{HGI}$ datasets (round 4 alpha and round 4 freeze). The color indicates the strength of linkage disequilibrium to the lead SNP rs35081325.

**Table S1: Hallmark pathways and their relevance scores**

| Hallmark pathway | Relevance score |
|---|---|
| INTERFERON_GAMMA_RESPONSE | 0.704 |
| ALLOGRAFT_REJECTION | 0.656 |
| INTERFERON_ALPHA_RESPONSE | 0.522 |
| MTORC1_SIGNALING | 0.434 |
| COMPLEMENT | 0.415 |
| HYPOXIA | 0.386 |
| P53_PATHWAY | 0.315 |
| TNFA_SIGNALING_VIA_NFKB | 0.263 |
| G2M_CHECKPOINT | 0.225 |
| XENOBIOTIC_METABOLISM | 0.217 |
| APOPTOSIS | 0.212 |
| ESTROGEN_RESPONSE_LATE | 0.200 |
| GLYCOLYSIS | 0.185 |
| ANDROGEN_RESPONSE | 0.178 |
| KRAS_SIGNALING_UP | 0.152 |
| ESTROGEN_RESPONSE_EARLY | 0.143 |
| MYC_TARGETS_V1 | 0.131 |
| IL2_STAT5_SIGNALING | 0.115 |
| UNFOLDED_PROTEIN_RESPONSE | 0.099 |
| UV_RESPONSE_UP | 0.081 |
| OXIDATIVE_PHOSPHORYLATION | 0.056 |
| EPITHELIAL_MESENCHYMAL_TRANSITION | 0.055 |
| ADIPOGENESIS | 0.045 |
| INFLAMMATORY_RESPONSE | 0.045 |
| DNA_REPAIR | 0.004 |
| CHOLESTEROL_HOMEOSTASIS | -0.002 |
| PROTEIN_SECRETION | -0.004 |
| MITOTIC_SPINDLE | -0.033 |
| MYC_TARGETS_V2 | -0.048 |
| FATTY_ACID_METABOLISM | -0.050 |
| APICAL_JUNCTION | -0.056 |
| MYOGENESIS | -0.058 |
| PI3K_AKT_MTOR_SIGNALING | -0.068 |
| IL6_JAK_STAT3_SIGNALING | -0.086 |
| PEROXISOME | -0.087 |
| KRAS_SIGNALING_DN | -0.113 |
| E2F_TARGETS | -0.120 |
| TGF_BETA_SIGNALING | -0.122 |

| | |
|---|---|
| SPERMATOGENESIS | -0.126 |
| UV_RESPONSE_DN | -0.128 |
| BILE_ACID_METABOLISM | -0.135 |
| REACTIVE_OXYGEN_SPECIES_PATHWAY | -0.141 |
| COAGULATION | -0.167 |
| HEME_METABOLISM | -0.202 |

**Table S2: Transcription factors and their relevance scores**

| Transcription factor | Relevance score |
|---|---|
| RELA | 0.530 |
| STAT1 | 0.380 |
| NFKB1 | 0.354 |
| MYCN | 0.285 |
| HDAC1 | 0.275 |
| USF1 | 0.267 |
| SP1 | 0.256 |
| PPARG | 0.209 |
| CIITA | 0.183 |
| TWIST2 | 0.108 |
| ETS1 | 0.094 |
| RFX5 | 0.094 |
| HIF1A | 0.092 |
| TWIST1 | 0.089 |
| EGR1 | 0.071 |
| MYC | 0.066 |
| USF2 | 0.065 |
| CEBPB | 0.056 |
| ATF4 | 0.055 |
| CREM | 0.024 |
| YY1 | 0.009 |
| FOS | 0.008 |
| JUN | 0.005 |
| IRF1 | 0.003 |
| XBP1 | -0.009 |
| SP3 | -0.024 |
| NR3C1 | -0.026 |
| TP53 | -0.026 |
| AR | -0.046 |
| E2F1 | -0.058 |
| CREB1 | -0.072 |

| | |
|---|---|
| EP300 | -0.073 |
| ESR1 | -0.077 |
| HSF1 | -0.091 |
| BRCA1 | -0.091 |
| RUNX1 | -0.098 |
| GATA1 | -0.099 |
| ATM | -0.116 |
| SPI1 | -0.123 |
| DNMT1 | -0.124 |
| TFAP2A | -0.138 |
| STAT3 | -0.156 |
| RUNX3 | -0.159 |
| SIRT1 | -0.182 |
| WT1 | -0.197 |
